# Intercellular Concentration Gradients of 3-Phosphoglycerate and Triose-Phosphate Demonstrate Operation of an Energy Shuttle in NAD-Malic Enzyme and Phospho*enol*ypyruvate Carboxykinase C_4_ Subtypes

**DOI:** 10.64898/2026.03.26.714472

**Authors:** Vittoria Clapero, Regina Feil, Stéphanie Arrivault, Mark Stitt

**Affiliations:** Max Planck Institute of Molecular Plant Physiology, Am Muehlenberg 1, 14476 Potsdam-Golm, Germany

**Keywords:** C_4_ photosynthesis, intercellular energy shuttle, leaf fractionation, NAD-malic enzyme, phospho*enol*pyruvate carboxykinase, 3-phosphoglycerate, triose phosphates

## Abstract

In C_4_ photosynthesis, incoming CO_2_ is incorporated in mesophyll cells (MC) into 4-carbon acids that diffuse to bundle sheath cells (BSC) and decarboxylated to generate a high CO_2_ concentration that suppresses the oxygenation reaction of Rubisco. Decarboxylation can occur by NADP-malic enzyme, (NADP-ME), NAD-malic enzyme (NAD-ME) or phospho*enol*pyruvate carboxykinase (PEPCK). NADP-ME generates NADPH in the BSC chloroplast and species that use it as the major route for decarboxylation typically have dimorphic BSC chloroplasts with little or no photosystem II. They operate an energy shuttle: much of the 3-phosphoglycerate formed in the Calvin-Benson cycle diffuses to the MC, enters the chloroplasts and is reduced to triose phosphates that return to the BSC. In species where carboxylation occurs mainly via NAD-ME or PEPCK, BSC chloroplasts possess photosystem II. Indirect evidence indicates they nevertheless have the capacity to operate an energy shuttle. We show here that NAD-ME and PEPCK species possess large pools of 3PGA and triose phosphates and, for two examples of each subtype, opposed concentration gradients of 3-phosphoglycerate and triose phosphates to drive rapid exchange between the BSC and MC. Reasons for and consequences of the widespread operation of the intercellular energy shuttle in C_4_ plants are discussed.

**Highlight Statement:** An intercellular energy shuttle in which 3-phosphoglycerate moves from the bundle sheath to the mesophyll and triose phosphates return to the bundle sheath is a general feature of C_4_ photosynthesis.

## Introduction

In C_4_ photosynthesis a biochemical CO_2_-concentrating mechanism (CCM) raises the CO_2_ concentration around Rubisco, suppressing the oxygenation reaction and the need for photorespiration (Hatch and Osmond, 1976; Hatch, 1987). This allows faster photosynthesis and increased water and nitrogen use efficiencies (von Caemmerer and Furbank, 2003). C_4_ photosynthesis evolved over 60 times in independent plant lineages 25-30 million years ago under falling atmospheric CO_2_ concentration (Christin *et al*., 2008; Sage *et al*., 2011). Although C_4_ species represent only ∼3% of current terrestrial plant species, they account for 23% of global primary production (Kellogg, 2013; Sage, 2016). C_4_ species have a specialized Kranz leaf anatomy with two distinct cell types: mesophyll cells (MC) and centrally-located bundle sheath cells (BSC). After entering the MC, CO_2_ is converted by carbonic anhydrase to bicarbonate that is assimilated by phospho*enol*pyruvate carboxylase (PEPC) to produce oxaloacetate (OAA). OAA is converted to malate or aspartate that move from the MC to the BSC, where they are decarboxylated to release CO_2_ and a C_3_ moiety, which moves back to the MC and is used to regenerate PEP. Intercellular movement occurs by diffusion, facilitated by the high plasmodesmata frequency at the MC-BSC interface (Osmond, 1971; Danila *et al*., 2016, 2018) and driven by large intercellular concentration gradients (Leegood 1985; Stitt and Heldt, 1985a, 1985b; Arrivault *et al*., 2017; Tonetti *et al*., 2025)

C_4_ species have conventionally been divided into three subtypes according to the main decarboxylating enzyme (Hatch, 1987): NADP-malic enzyme (NADP-ME) in the plastid, NAD-malic enzyme (NAD-ME) in the mitochondria or phospho*enol*pyruvate carboxykinase (PEPCK) in the cytosol (Figures 1A, B and C). This in turn influences which C_4_ acids and C_3_ moieties move between the MC and the BSC, which intracellular transport steps are required as well as energy requirements in the MC and BSC (Supplementary Figure S1, Supplementary Table S1). In the NADP-ME subtype, OAA is reduced by plastidic NADP-malate dehydrogenase to malate which diffuses to the BSC, enters the chloroplast and is decarboxylated by NADP-ME, releasing CO_2_, NADPH and pyruvate. Pyruvate returns to the MC and used by pyruvate, phosphate dikinase (PPDK) to regenerate PEP. In the NAD-ME subtype, OAA is transaminated in the MC cytosol by aspartate aminotransferase (AspAT) to produce aspartate which moves to the BSC, enters the mitochondria and is converted by AspAT and NAD-malate dehydrogenase to malate that is decarboxylated by NAD-ME, releasing CO_2_, NADH and pyruvate. The pyruvate is converted by alanine aminotransferase (AlaAT) to alanine, which returns to the MC and deaminated to produce pyruvate that is used by PPDK to regenerate PEP. In the PEPCK subtype, both malate and aspartate are produced from OAA in the MC. The aspartate diffuses to the BSC cytosol, where it is deaminated by AspAT before being decarboxylated by PEPCK, releasing CO_2_ and PEP. The malate moves to the BSC where it is decarboxylated by NAD-ME or sometimes NADP-ME.

**Figure 1.**
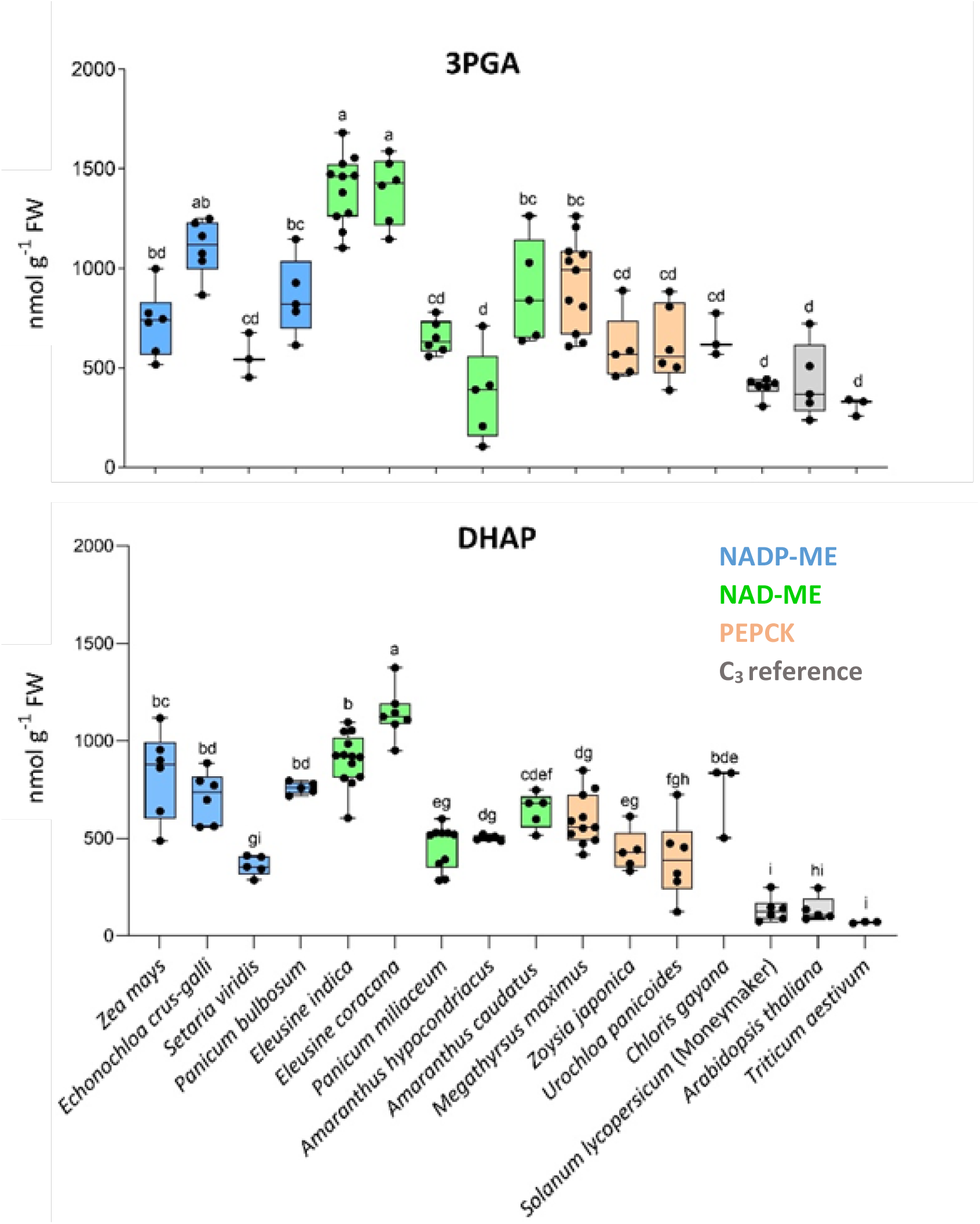
Overall content of 3PGA and DHAP in the C_4_ species panel and three reference C_3_ species. Blue = C_4_ NADP-ME species, green = C_4_ NAD-ME species, orange = C_4_ PEPCK species, grey = C_3_ species. The boxes represent the lower and upper quartile (25%), the horizontal line in each box indicates the median and the whiskers represent minimum and maximum values. The letters on the graph indicate significant differences between the means of each species, with p-value < 0.05 (ANOVA and Tukey’s test performed in GraphPad Prism ver 10.2.2). Plants were grown in a 14h light (28°C)/10h night (20°C) regime at an irradiance of 590 μmol photons m^-2^ s^-1^ and 80% RH. A section of around 10cm from the central part (i.e., excluding the tip) of the third fully expanded leaf was harvested in the light field and quickly quenched in liquid nitrogen without shading and while keeping it horizontal to the light. The 3PGA and DHAP content and number of independent replicates are provided in Supplementary Table S3. Data for *Triticum aestivum* from Arrivault *et al*. (2017).

Despite their apparent clarity, textbook diagrams of C_4_ photosynthesis represent simplifications. It is widely accepted that most C_4_ plants operate more than one decarboxylation pathway (Furbank, 2011; Wang *et al*., 2014, von Caemmerer and Furbank, 2016; Weissmann *et al*., 2016; Arrivault *et al*., 2017; Baccolini *et al*., 2026). This may increase flexibility, aid adjustment to different conditions, and decrease the size of the concentration gradients required to drive intercellular movement of individual metabolites. There are also open questions regarding the pathways themselves.

One concerns the extent to which there is transfer of energy between the MC and BSC. This is essential in NADP-ME subtypes because they have dimorphic chloroplasts in the BSC with little or no photosystem II (PSII) activity (Hatch 1987; Friso *et al*., 2010). They operate an energy shuttle, in which 3-phosphoglycerate (3PGA) diffuses from the BSC to the MC where it is reduced to triose phosphates (triose-P) that diffuse back to the BSC. Based on idealised pathway stoichiometry, each molecule of CO_2_ produced by NADP-ME leads to the formation of two molecules of 3PGA. One can be reduced using the molecule of NADPH produced by NADP-ME and the other moves to the MC. The actual stoichiometry depends on how far PSII function is abrogated in the BSC chloroplast and the extent of back-leakage of CO_2_. Evidence for operation of this energy cycle in NADP-ME species includes high activities of phosphoglycerate kinase (PGK), triose phosphate isomerase (TPI) and NADP-glyceraldehyde-3-phosphate dehydrogenase (NADP-GAPDH) in MC chloroplasts (Raghavendra and Das, 1978), expression of the triose phosphate:phosphate translocator (TPT) that facilitates exchange of 3PGA and triose-P across the chloroplast envelope membrane in the MC (Bräutigam *et al*., 2008; Chang *et al*., 2012; Emms *et al*., 2016; Mattinson and Kelly, 2025). There is also strong functional evidence, including high rates of 3PGA reduction in isolated MC chloroplasts (Furbank *et al*., 1983), the presence of large overall pools of 3PGA and triose P compared to other Calvin-Benson cycle (CBC) intermediates (Leegood and von Caemmerer, 1989; Arrivault *et al*., 2019) and, especially, experimental demonstration of a large concentration gradient of 3PGA between the BSC and MC and a large opposing gradient of triose-P between the MC and the BSC in *Zea mays* (Leegood 1985; Stitt and Heldt, 1985a, 1985b; Arrivault *et al*., 2017) and *Setaria viridis* (Tonetti *et al*., 2025). These gradients drive rapid diffusion between the two cell types.

It is less clear whether an analogous energy shuttle operates in NAD-ME and PEPCK subtypes, which have fully operational PSII in their BSC chloroplasts. Theoretical considerations including the relatively low requirement for ATP and reducing power in the MC of NAD-ME and PEPCK subtypes (Supplementary Table S1) indicate the shuttle may be needed to utilise surplus energy in the MC. Indeed, its operation was already assumed by Hatch (1987) (see also Edwards and Voznesenskaya, 2011; Brautigam *et al*., 2014; von Caemmerer and Furbank, 2016). Scattered experimental evidence indicates that NAD-ME and PEPCK species have the capacity to operate an energy shuttle. This includes the presence and/or expression of PGK, TPI and NADP-GAPDH in the MC chloroplast from the NAD-ME species *Amaranthus paniculatus* (Raghavendra and Das, 1978), *Eleusine indica, Panicum miliaceae* (Ku and Edwards, 1975) and *Panicum virgatum* (John *et al*., 2014) and the PEPCK species *Panicum texanum* and *Urochloa panicoides* (Ku and Edwards, 1975; see also discussion in Bräutigam *et al*., 2014), the expression of TPT in the MC chloroplast of the NAD-ME species *P. virgatum* and *G. gynandra* (Aubry *et al*., 2014; Rao *et al*., 2016), rapid 3PGA reduction by isolated MC chloroplasts from the NAD-ME species *P. miliaceum* and the PEPCK species *Megathyrsus maximus* (Furbank *et al*., 1983) and a report of high 3PGA and triose-P levels in the NAD-ME species *Amaranthus edulis/caudatus* (von Leegood and Caemmerer, 1988). However, to our knowledge, there has not been a broad survey of 3PGA and triose-P levels or an experimental demonstration of intercellular gradients of 3PGA and triose P in NAD-ME and PEPCK species.

Whether an energy shuttle operates in NAD-ME and PEPCK species is not only important to fully understand the pathways of carbon and energy flow in these subtypes. It is also relevant for understanding why most NADP-ME species have dimorphic BSC chloroplasts with abrogated PSII. One explanation would be that the NADP-ME reaction provides NADPH, allowing partial or complete loss of PSII, and that the energy shuttle evolved as a response to loss of PSII in the BSC chloroplast. An alternative explanation would be that the energy shuttle is a widespread feature of C_4_ photosynthesis, irrespective of the decarboxylation pathway, and the abrogation of PSII in BSC chloroplasts of NADP-ME species is a specific adaption to allow efficient operation of NADP-ME (Bräutigam *et al*., 2018).

A second open question relates to the route by which, the PEP moves from the BSC to the MC in PEPCK species. Various routes have been considered: that PEP itself moves (Rathnam and Edwards, 1977), that PEP is converted by pyruvate kinase to pyruvate that is transaminated and returns as alanine (Hatch and Kagawa 1976; Hatch, 1979), or that PEP is converted in the BSC by enolase and phosphoglycerate mutase (PGM) to 3PGA that moves to the MC where PGM and enolase convert it back to PEP (Huber and Edwards, 1975). There is little experimental evidence for any of these routes. Movement of PEP itself would require a concentration gradient between the BSC and MC, but there are no data available on the intercellular distribution of PEP in PEPCK species. For the other two routes, it is unclear if there is sufficient enzymic capacity. Whilst the PEPCK species *Spartina anglica* Hubb. L has high in vitro activity of alanine aminotransferase, measured in vitro activities of pyruvate kinase and PPDK were insufficient to sustain the measured rate of photosynthesis (Smith *et al*., 1982; Smith and Woolhouse, 1983). Further, transcriptome studies revealed weaker expression of PPDK in the PEPCK species *M. maximus* than the NAD-ME species *Cleome gynandra* (Bräutigam *et al*., 2014). The hypothesis that PEP is converted to 3PGA before moving to the MC is thermodynamically attractive; the enolase and PGM reactions are typically near to equilibrium *in vivo*, and their combined equilibrium constant would lead to a 3PGA concentration that is about three-fold higher than the PEP concentration (Newsholme and Start, 1973; Leegood and von Caemmerer, 1989), amplifying the concentration gradient to drive diffusion to the MC. However, the *in vitro* activities of enolase and PGM in *S. anglica* Hubb. were insufficient to sustain the measured rate of photosynthesis L (Smith *et al*., 1982; Smith and Woolhouse, 1983), and in vitro activities of PGM and enolase in the PEPCK *species P. texanum* and *U. panicoides* were no higher than in other C_4_ subtypes or the C_3_ reference species *Triticum aestivum* (Ku and Edwards, 1975).

The main aim of the following study was to experimentally clarify if an energy shuttle operates in NAD-ME and PEPCK subtypes. We first asked if NAD-ME and PEPCK species contain large pools of 3PGA and triose-P, comparable to those in NADP-ME species. This would be a prerequisite to generate large intercellular concentration gradients. We then adapted a method developed to partially enrich MC and BSC material from *Z. mays* leaves by differential homogenisation and filtration through nylon nets of different apertures, with all steps being performed in liquid N_2_ to preserve metabolite levels. (Stitt and Heldt, 1985a, 1985b). This allowed us to analyse the intercellular distribution of 3PGA and triose-P in two NAD-ME species and two PEPCK species. A minor aim was to ask if there is an appreciable concentration gradient for PEP between the BSC and MC in PEPCK species.

## Materials and methods

### Plant material, growth and harvest

Plant species and their origin are listed in Supplementary Table S2 (see also Supplementary Figure S2). Both for comparison of overall metabolite levels and for analysis of metabolite distribution, plants were grown in repeated batches, subject to availability of seeds and requirements for experiments, but always using the same substrate and climatic conditions. Seeds were placed on 2% v/v sucrose agar and germinated in 8h light (25°C)/16h night (25°C) cycle with irradiance of 110 μmol photons m^-2^ s^-1^ for 2 weeks. The plants were then transferred to soil and grown at 14h light (28°C)/10h night (20°C), irradiance of 590 μmol photons m^-2^ s^-1^ and 80% RH. A section of around 10 cm from the central part (i.e., excluding the tip) of the third fully expanded leaf was harvested in the light field and quickly quenched in liquid nitrogen keeping it horizontal to the light and without shading.

### Enzyme activity measurement

Phosphoribulokinase (PRK) activity was measured, in a continuous assay modified from Gardemann *et al*. (1983) with 10 mM MgCl_2_, 100 mM Tricine/KOH (pH 8), 50 mM KCl, 1mM ATP, 0.2 mM NADH, 3 mM PEP, 0.3 units pyruvate kinase, 1 unit lactate dehydrogenase, starting the reaction by adding 5 mM ribose-5 phosphate (R5P) and 0.02 units phosphoriboisomerase (PRI), in a final volume of 210μl. Absorption at 340 m was measured using an Agilent BioTek ELX808 94 well plate reader (Agilent Technologies, Santa Clara, CA, USA). R5P was chosen as starting substrate instead of Ru5P for cost reasons, and PRI was added to supplement endogenous enzyme in the complete conversion of R5P to ribulose-5 phosphate (Ru5P). The assay contained 0.025 mg FW material for *Zea mays, Eleusine indica* and *Panicum miliaceum* and 0.01 mg of FW material for *Megathyrsus maximus* and *Urochloa panicoides*. In preliminary experiments, the added amounts of coupling enzymes were optimised to minimise extract-independent oxidation of NADH by trace contaminants in the coupling enzymes. The chosen concentration of phosphoriboisomerase allowed rapid conversion of R5P to Ru5P, so that the PRK reaction started without appreciable lag after addition of R5P and PRI. The measured activity was corrected for the blank given by the coupling enzyme PRI in the complete reaction minus the soluble enzyme extract. Preliminary checks also showed that the chosen concentrations of pentose-P and ATP were saturating for PRK activity, and of NADH for the reaction with the coupling enzyme malate dehydrogenase (MDH). Each assay was carried out with two technical replicates.

PEPC activity was measured in a continuous assay modified from Meyer at al. 1988) in 5 mM MgCl_2_, 25 mM Tris/HCl (pH 8), 2 mM DTT, 1 mM KHCO_3_, 5mM glucose-6 phosphate (G6P), 0.2 mM NADH, 2 units of MDH, started by adding 5 mM PEP, in a final volume of 210μl. Absorption at 340 m was measured on an Agilent BioTek ELX808 94 well plate reader. The assay contained 0.05 mg FW material for *Z. mays, E. indica* and *P. miliaceum* and 0.01 mg FW material for *M. maximus* and *U. panicoides*. Preliminary checks showed that the coupling enzymes and concentrations of PEP, KHCO_3_ and the activator G6P were saturating for PEPC activity. Each assay was carried out with two technical replicates.

NADP-ME was measured, modified from Hatch and Mau (1977a), in a continuous assay at 340 nm based on the production of NADPH in 25 mM Tricine/KOH (pH 8.3), 200 μM EDTA, 5 mM malate, 500 μM NADP, started by adding 2mM MgCl_2_ in a final volume of 210μl. The assay contained protein extract from 0.1 mg of FW material for all species assayed. Absorption at 340 m was measured in an Agilent BioTek ELX808 94 well plate reader. NAD-ME activity measured, modified from Hatch *et al*. (1982), in a continuous assay in 25 mM HEPES/KOH (pH 7.5), 200 μM EDTA, 5 mM DTT, 5 mM malate, 2 mM NAD, 4mM MnCl_2_, starting the assay by 5 mM coenzyme A (CoA) in a final volume of 210μl. The assay contained 0.1 mg of FW material for all species assayed. Absorption at 340 m was measured in an Agilent BioTek ELX808 94 well plate reader. PEPCK activity was measured, modified from Hatch and Mau (1977b), in a continuous spectrophotometric assay following the disappearance of OAA at 280 nm in 50 mM HEPES/KOH (pH 7.5), 10 mM MgCl_2_, 100 mM KCl, 1 mM MnCl_2_, 15 mM DTT, 300 μM OAA, 1 unit of pyruvate kinase, starting the reaction by adding 10μl of a stock solution to generate 200 μM ATP in a final volume of 895μl (quartz cuvette). The assay contained 2 mg FW for all species. Absorption at 280 nm was measured on a Shimadzu UV-2600 UV-spectrophotometer (Shimadzu, Kyoto, Japan).

### Metabolite determination

An aliquot of 50 mg frozen material was first treated with 400 μl 16% w/v trichloroacetic acid (TCA) in diethylether. After incubating for 20 minutes on ice, 200 μl 16% TCA, 5 mM EGTA in water was added and mixed. After a 2 hours long further incubation on ice, the samples were centrifuged for 5 minutes at 14000 rpm in a pre-cooled centrifuge. The water phase is then isolated, washed four times with 400 μl water saturated diethylether and eventually neutralised with 1M KOH/200 mM TEA if needed (adapted from Jelitto *et al*., 1992). Prior to measurement, the extract was treated with activated charcoal and spun down at 14000 rpm in a pre-cooled centrifuge to remove aromatic compounds and other compounds absorbing in the UV spectrum.

3PGA was enzymatically assayed in a final volume of 910 μl containing 100 mM Tris/HCl (pH 8.1), 5 mM MgCl_2_, 0.03 mM NADH and 0.06 mM ATP starting the reaction by adding 0.5 units of GAPDH and 0.2 units of PGK. DHAP was enzymatically assayed in in a final volume of 150 μl containing 100 mM Tris/HCl (pH 8.1), 5 mM MgCl_2_, 0.2 mM NADH, starting the reaction by adding 0.1 units of GDH. PEP was enzymatically assayed in a final volume of 910 μl with 100 mM Tris/HCl (pH 7.5), 5 mM MgCl_2_, 0.1 mM NADH, 0.9 mM ADP, 2 units of LDH, starting the reaction by adding 2 units of PK. NADH extinction was measured at 340 nm using a Shimadzu UV-2600 UV-spectrophotometer.

Fructose-1, 6 bisphosphate (FBP) and fructose-6 phosphate (F6P) were determined in the filtered aqueous phase of a chloroform/methanol extraction by high-performance anion-exchange liquid chromatography coupled to tandem mass spectrometry after Lunn *et al*. (2006) modified as in Figueroa *et al*. (2016)). Aliquots (25 μL) of the filtered extracts were spiked with 2.5 pmol [U-^13^C]F6P and 2.5 pmol [U-^13^C]FBP (25 μL) before injection onto the column to allow correction for ion suppression and other matrix effects.

### Leaf fractionation to obtain material enriched in MC and BSC

Leaf fractionation followed Stitt and Heldt (1985a) and Stitt *et al*. (1989), with modifications. About 1 g FW deep-frozen leaf material was partially homogenized in liquid N_2_ in a ceramic mortar and pestle that was pre-chilled with liquid N_2_. The extent of homogenisation was established by empirical trial and assessed by visual inspection. The resulting powder was resuspended in excess liquid N_2_ and filtered successively through 200 μm, 80 μm and 40 μm nylon nets. For each nylon net, flow-through was collected in a chilled mortar and material retained on the nets was washed with extra liquid N_2_ to ensure thorough filtration. The combined flow-through was then filtered through the next nylon net. The protocol generated four fractions: residues on the 200 μm, 80 μm and 40 μm nylon nets, and the final flow-through (termed fractions 1, 2, 3 and 4, respectively, Supplementary Figure S3A). Leaf material was at liquid N_2_ temperatures at all times and eventually scraped into pre-frozen Eppendorf tubes that were stored at -70°C until further analysis. For each species, many fractionations were performed with separate batches of leaf material; each fractionation is termed an experiment.

Weighed subaliquots were extracted for measurement of marker enzyme activities and metabolite content (see above). Given the fibrous composition of fractions 1 and 2, an additional homogenization step was added using an oscillating ball mill (Retsch) using two 30 Hz 1 min pulses (while frozen).

### Mathematical extrapolation of metabolite distribution between MC and BSC

Metabolite distribution between the MC and BSC was estimated based on Gerhardt and Heldt (1984) and Stitt and Heldt (1985a), with modifications to allow fuller use of the dataset and statistical evaluation of the quality of the estimates in each individual experiment. The activities of enzymes (μmol g^-1^ FW min^-1^) and the amounts of metabolites (nmol g^1^-FW) were first summed across all four fractions, and the value in each fraction expressed as a percentage of the total. This transformation generates values for enzyme activities and metabolite amounts that are dimensionless and thus directly comparable. Assuming the ratio between the amount of the metabolite deriving from the MC (X _mesophyll_) and from the BSC (X _bundle sheath_) and the activities of the corresponding marker enzymes PEPC and PRK is constant across all fractions:

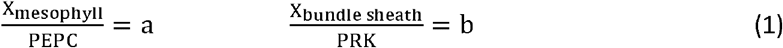

Assuming the metabolite is distributed only between the MC and BSC, the total amount quantified in a fraction consists of the MC and BSC portions:

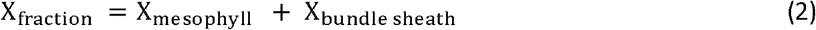

Substitution of equation (1) in (2) gives:

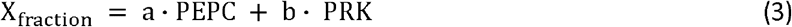

Theoretically, coefficients *a* and *b* can take values between 0 and 1, and represent the contribution of the MC and BSC respectively to the metabolite content of a given fraction. Equation (3) can be reorganized as a linear function, with the form:

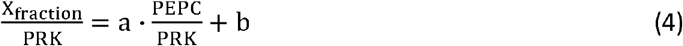

By plotting the values of 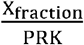 and 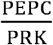 (y- and x-axis, respectively) for the four fractions and performing a regression, *a* and *b* can be estimated (slope and y-intercept, respectively).

A second linear function can be derived in an analogous manner:

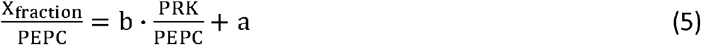

Plotting the values of 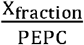 and 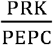 (y-and x-axis, respectively) for the four fractions and performing a regression allows estimation of *a* and *b* (in this case, y-intercept and slope, respectively).

Illustrative examples of the plots are provided in Supplementary Figure S3B. As the transformed values for enzyme activities and metabolite amounts are dimensionless (see above), metabolite distribution can be read directly from the plots. The proportion of metabolite X in the MC is given by the slope of the left-hand plot and the y-intercept of right-hand plot (in this example, 0.75 and 0.86, respectively), and the proportion of metabolite X in the BSC is given by the y-intercept of the right-hand plot and the slope of the right-hand plot (in this example, 0.26 and 0.35, respectively). In principle, the estimates from both plots should be the same, and the proportions estimated for the BSC and MC should sum to one. Due to experimental noise this will not be precisely so, and inspection of the values provides a qualitative check on the reliability of the estimates for a given experiment.

Goodness of fit of a linear regression is usually assessed by calculating Pearson’s coefficient (R^2^). However, when the slope was very small (i.e., the line was near-horizontal) R^2^ was often low even when all points were close to the line (not shown). This follows from the intrinsic null hypothesis of the Pearson’s test, viz. that there is no linear function of x which can explain the variation of *y* better than the null hypothesis y=k (a perfectly horizontal line). For this reason, instead of calculating R^2^, p-values were generated for the slope and for the y-intercept using the *lm*() function of RStudio software version 4.3.1. The null-hypothesis is that the predictor (slope or intercept) is equal to zero, so that it has no effect on describing variation of the dependent variable *y*. A low p-value on either slope or intercept indicates the predictor is a meaningful addition to the model (viz. the linear regression).

Significance of the difference of the linear regressions for 3PGA and DHAP (per species, either based on Equation 4 or Equation 5) was tested in RStudio (software version 4.3.1) using a linear model with an interaction term (Equation 6):

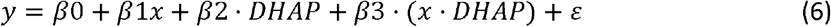

This model takes the levels of 3PGA as the reference levels and tests for β_0_ = intercept for 3PGA, β_1_ = slope for 3PGA, β_2_ = difference in intercept between DHAP and 3PGA, β_3_ = difference in slope between DHAP and 3PGA. Significant p-values associated with the coefficients β_2_ and β_3_ indicate (respectively) differences in the intercepts and slopes of the linear regressions for 3PGA and DHAP.

## Results

### Overall content of 3PGA and triose-P in a panel of NADP-ME, NAD-ME and PEPCK subtypes

A panel of four NADP-ME species (*Zea mays, Setaria viridis, Echinochloa crus-galli, Panicum bulbosum*), five NAD-ME species (*Eleusine indica 81049, Eleusine coracana, Panicum miliaceum, Amaranthus caudatus, Amaranthus hypochondriacus*) and four PEPCK species (*Megathyrsus maximus 78380, Zoysia japonica, Urochloa panicoides, Chloris gayana*) was assembled (Supplementary Table S2), mainly from the Poaceae but with two eudicot NAD-ME species from the Caryophyllales (Supplementary Figure S2). 3PGA and DHAP levels were measured in mature leaves of these and two reference C_3_ species (*Arabidopsis thaliana, Solanum lycopersicum, Triticum aestivum*).

Except for the NAD-ME eudicot *A. hypochondriacus*, the levels of 3PGA in the NAD-ME and PEPCK species were higher than in the C_3_ reference species and comparable if not higher than in the NADP-ME species. *Eleusine indica* and *Eleusine coracana* had almost double the amount in the model C_4_ species *Z. mays* (average 1396 nmol g^-1^ FW and 1392 nmol g^-1^ FW compared to 724 nmol g^-1^ FW, respectively). *A. hypochondriacus* had levels comparable to *A. thaliana* and *S. lycopersicum*.

Levels of DHAP in the NAD-ME and PEPCK species were at least three-fold higher than the reference C_3_ species and comparable (range 455 nmol g^-1^ FW to 1140 nmol g^-1^ FW) to the model NADP-ME species *Z. mays* (average 826 nmol g^-1^ FW). *E. indica* and *E*.*e coracana* again had the highest DHAP levels (905 nmol g^-1^ FW and 1140 nmol g^-1^ FW, respectively). The C_4_ species with the lowest levels of DHAP was the NADP-ME species *S. viridis* (360 nmol g^-1^ FW) but this was still almost three-fold higher than in the two reference C_3_ species (around 133 nmol g^-1^ FW in both).

Levels of F6P and FBP in the NAD-ME and PEPCK species were in the same range as in NADP-ME species and the C_3_ reference species (Supplementary Figure S4; Supplementary Table S4). Thus, the high levels of 3PGA and DHAP in the NAD-ME and PEPCK species was not due to generally higher levels of CBC intermediates.

Two NAD-ME species (*P. miliaceae, E. indica*) and two PEPCK species (U. *panicoides, M. maximus*) were chosen to perform leaf fractionation experiments. To check that the dominant decarboxylation route in these species under our growth conditions corresponded to that described in the literature, NADP-ME, NAD-ME and PEPCK activity was measured in these four species, together with *Z. mays* as a reference NADP-ME species (Supplementary Figure S5). The two NAD-ME species had high NAD-ME activity and detectable but much lower NADP-ME activity. The two PEPCK species had high PEPCK activity, much lower NADP-ME activity and very low NAD-ME activity.

### Intercellular distribution of 3PGA and triose-P in selected NAD-ME and PEPCK species

Leaf material was processed to produce fractions enriched in BSC and MC using a method described in Stitt and Heldt (1985a). Briefly, leaf material was partially homogenised in liquid N_2_ using a mortar and pestle, and sequentially filtered in liquid N_2_ through a 200 µm nylon net, an 80 µm nylon net and a 40 µm nylon net (Supplementary Figure S3, for more details see Methods). Due to the higher physical robustness of the vascular tissue with their bundle sheath strands, this procedure generates four fractions; material retained in the 200 µm net (F1, corresponding mainly to large fragments that are not enriched for cell type), material retained in the 80 µm net (F2, enriched for BSC), material retained on the 40 µm net (F3, not strongly enriched for cell type) and material that passed through the 40 µm net (F4, enriched for MC). The extent of enrichment was assessed by measuring the activity of MC and BSC enzyme markers, phospho*enol*pyruvate carboxylase (PEPC) and phosphoribulokinase (PRK) respectively, in each fraction.

The distribution of a given marker enzyme was calculated by expressing the activity in each fraction as a percentage of the summed activity in all four fractions. The extent of enrichment in each fraction is then given by comparing the distribution of the two marker enzymes. Similarly, the distribution of a given metabolite was calculated by expressing the amount in each fraction as percentage of the summed amount in all four fractions. If the metabolite is exclusively in the BSC then its distribution will resemble that of PRK, if it is exclusively in the MC its distribution will resemble that if PEPC, and if it is located in both cell types then it will show an intermediate distribution. The distribution of the metabolite can be graphically solved by plotting %metabolite/%PRK vs %PEPC/%PRK or %metabolite/%PEPC vs %PRK/%PEPC where %metabolite, %PEPC and %PRK are the percentage of the total metabolite, total PRK and total PEPC that is present in a given fraction. The proportion of metabolite in the MC is given by the slope of the first plot and the y-axis intercept of the second plot, and the proportion of metabolite in the BSC is given by the y-axis intercept of the first plot and the slope of the second plot (for more details, see Materials and Methods and Supplementary Figure S3). Theoretically, the two estimates should be identical and the proportion in the MC and BSC should sum to unity. However, there may be some divergence due to experimental noise. Further, the approach assumes that all of the PEPC is in the MC, all of the PRK is in the BSC and the metabolite is restricted to the MC and BSC, which may not be completely the case.

For each species, multiple experiments were performed. ‘Experiment’ refers to the group of four fractions obtained from differential filtration of one batch of leaf material. Each experiment was assigned a unique name given by a species label (E= *E. indica*, P= *P. miliaceum*, T= *M. maximus*, U= *U. panicoides*). Some experiments were rejected because there was poor enrichment. Supplementary Table S5 summarizes 4-5 successful experiments for each species, giving the estimated proportion of 3PGA or DHAP in the MC and in the BSC. Two estimates and corresponding p-values are given; one from plots based on equation 4 and one from plots based on equation 5 (see Materials and Methods). Similar estimates were usually obtained from both plots, the estimated distributions were usually significant when estimated from the intercept and often significant when estimated from the slope and, in all experiments, the estimated proportion in the MC and BSC summed to close to unity. In a few individual experiments, the proportion of a metabolite assigned to one cell type was >1, with a negative value for the other cell type. This may reflect experimental noise or location of a metabolite in other cell types in addition to the BSC and MC (see Discussion).

For further data analyses, the data points of all experiments were combined and used for regression analyses (Supplementary Figure S6, Supplementary Table S5). Similar results were obtained from plots based on equation 4 and equation 5, almost all regressions were significant and the proportion in the BSC and MC again summed to close to unity These values are summarised in Figure 2. In two cases where regression analysis indicted a negative content (DHAP in the BSC in *P. miliaceum*,3PGA in the MC in *U. panicoides*) the value was reset as zero. A larger proportion of the 3PGA was assigned to the BSC than the MC in all four species. A larger proportion of the DHAP was assigned to the MC than the BSC in *E. indica, P. miliaceum* and *U. panicoides*, but not *M. maximus*. Furthermore, regressions performed on 3PGA were significantly different from regressions performed on DHAP for all species when the regression were performed with equation 4, and for *P. miliaceum* and *U. panicoides* when the analyses were performed with equation 5 (Supplementary Figure S6).

**Figure 2.**
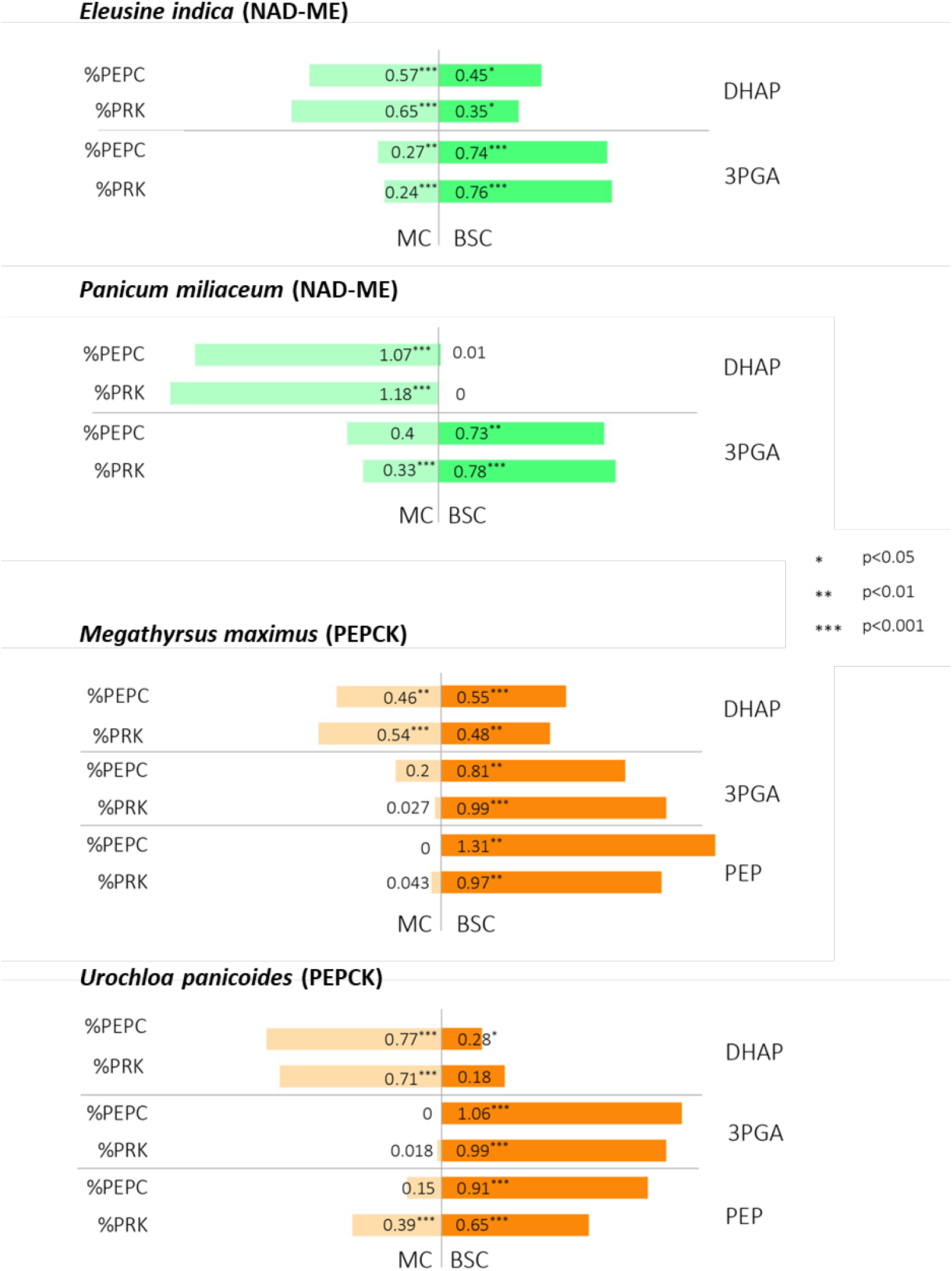
Summary of estimated percentage of 3PGA, DHAP and PEP in the MC and BSC of *Eleusine indica, Panicum miliaceum, Megathyrsus maximus* and *Urochloa panicoides*. The data and calculations are provided in Supplementary Dataset S1 and are summarized in Supplementary Table S4. The estimates are those from a combined analysis of all experiments performed for a given species. Two estimates are given for each species, based either on data normalized on PEPC (% PEPC) or on data normalized on PRK (%PRK) (for details see Methods and Supplementary Text). The estimated proportions are given as a fraction of the total metabolite in the leaf material (total = 1). The significance values of each estimate (tested as its significance in the linear regression describing the metabolite’s distribution, see Material and Methods) is indicated, following the legend.

To estimate amounts in the BSC and MC, the proportions in Figure 2 were multiplied by the total amounts from the experiment of Figure 1. The results are summarised in Table 1. There were much larger amounts of 3PGA in the BSC than the MC for all four species (254-729 nmol g^-1^ FW difference) and there were much larger amounts of DHAP in the MC than the BSC for *E. indica, P. miliaceum* and *U. panicoides* (81-455 nmol g^-1^ FW difference). ‘Negligible’ indicates the two cases where the regression analysis indicted a negative content (DHAP in the BSC of *P. miliaceum*, 3PGA in the MC of *U. panicoides*). The 3PGA/DHAP ratio was much higher in the BSC than the MC (Supplementary Table S6; ratios based for DHAP in the BSC in *P. miliaceum* are given as ‘HIGH’ and ratios based on 3PGA in the MC in *U. panicoides* as ‘LOW’).

**Table 1.** Summary of the estimated amounts in the MC and BSC. The average total content of 3PGA and DHAP (Figure 1) was assigned to the MC and BSC based on the proportional distribution given in Figure 2, using the average of the value estimated based on equation 4 and equation 5 (see Methods and Supplementary Table S5). In two cases where the regression analysis indicted a negative content (DHAP in the BSC in *P. miliaceum*, 3PGA in the MC in *U. panicoides*) the amount is indicated as negligible (negl). ‘Difference’ gives the difference between the amounts in the BSC and MC for 3PGA, and the difference between the amounts in the MC and BSC for DHAP (content is set as zero for 3PGA in the MC in *U. panicoides* and DHAP in the BSC in *P. miliaceum*). The difference for DHAP content in *M. maximus* has a negative sign, as the average of the estimates from equation 4 and equation 5 indicates slightly larger amount of DHAP in the BSC than the MC.

|  | Amount (nmol g<sup>-1</sup> FW) |  |  |
| --- | --- | --- | --- |
|  | MC | BSC | Difference |
| ***. indica (NAD-ME)*** |  |  |  |
| 3PGA | 355 | 1047 | Δ692 |
| DHAP | 498 | 416 | Δ81 |
| ***P. miliaceum (NAD-ME)*** |  |  |  |
| 3PGA | 237 | 492 | Δ254 |
| DHAP | 455 | negl | Δ455 |
| ***M. maximus (PEPCK)*** |  |  |  |
| 3PGA | 105 | 834 | Δ729 |
| DHAP | 306 | 297 | (-Δ8) |
| PEP | negl | 210 | Δ210 |
| ***U. panicoides (PEPCK)*** |  |  |  |
| 3PGA | negl | 615 | Δ615 |
| DHAP | 292 | 91 | Δ201 |
| PEP | 66 | 191 | Δ125 |

Amounts were converted to concentrations based on the reported volume of the cytoplasm of the MCs and BSCs in *Z. mays* leaves (Osmond. 1971; Hattersley, 1984; Furbank and Hatch, 1987; Jenkins *et al*., 1989; see legend of Table 2). The estimated concentrations are approximations because the volumes of the BSC and MC in the NAD-ME and PEPCK species may different from those in *Z. mays*. That said, they reveal substantial concentration gradients of 3PGA from the BSC to the MC in all four species, and substantial concentration gradients of DHAP from the MC to the BSC in for *E. indica, P. miliaceum* and *U. panicoides*, and a small gradient of DHAP in *M. maximus*.

**Table 2.** Estimated intercellular concentration gradients. Concentrations were calculated based on the internal leaf structure data available for *Z. mays* where BSCs occupy 19% of the leaf and the cytoplasm occupies around 50% of BSC volume (Furbank and Hatch, 1987; Jenkins *et al*., 1989), while MCs occupy ∼3.8 times of the leaf (Hattersley, 1984) and their cytoplasm occupies 10% of cell volume (Osmond, 1971) (see text). The metabolite contents (nmol/g FW) from Table 1 were divided by 9.5 (0.19*0.5*1000) to obtain the BSC concentration and by 7.22(0.19*3.8*0.1*1000) factor to obtain the MSC concentration. Concentration is set as zero for 3PGA in the MC in *U. panicoides* and for DHAP in the BSC in *P. miliaceum*. For *M. maximus* a small DHAP concentration gradient from MC to BSC is present, as the estimated cytoplasmic volume is smaller for MC than BSC.

| Metabolite | Gradient direction | Species |  |  |  |
| --- | --- | --- | --- | --- | --- |
|  |  | NAD-ME |  | PEPCK |  |
|  |  | *Eleusine indica* | *Panicum miliaceum* | *Megathyrsus maximus* | *Urochloa panicoides* |
| 3PGA | BSC → MC | 6.1 mM | 1.9 mM | 7.3 mM | 6.4 mM |
| DHAP | MC → BSC | 2.5 mM | 6.3 mM | 0.9mM | 3.1 mM |
| PEP | BSC → MC | - | - | 2.2 mM | 1.1 mM |

### Intercellular distribution of PEP in PEPCK species

In PEPCK species, it is a matter of debate how PEP moves from the BSC to the MC (see Introduction). One possibility is that some may return as PEP. We used the material from the above experiments to ask if there is a concentration gradient for PEP between the BSC and the MC. The overall PEP content in *M. maximus* and *U. panicoides* was 210 (n=5, SD= 58) and in 245 nmol g^-1^ FW (n=4, SD= 62), respectively. The estimated proportion in the BSC and MC is shown for analyses of the individual experiments and of the combined data set in Supplementary Table S7 and summarised in Figure 2, and estimated concentration gradients are given in Table 2. In *M. maximus*, PEP was almost exclusively located in the BSC, with an estimated concentration gradient of 2.2 mM, about 30% of that of 3PGA. In *U. panicoides*, PEP amounts were higher in the BSC than the MC, but with a smaller estimated concentration gradient of 1.1mM, <20% that of 3PGA.

Conversion of PEP to 3PGA in the BSC and conversion of 3PGA back to PEP in the BSC would require considerable activities of PGM and enolase in both cell types. Activities were investigated in four PEPCK species, two NAD-ME species, two NADP-ME species and two C_3_ species (Supplementary Figure S7). Enolase and PGM activities in PEPCK species (0.6-1.2 and 4.4-9.5 µmol g^-1^ FW min^-1^, respectively) resembled those previously reported for the PEPCK species *U. panicoides* (2.6 and 3.7 µmol mg^-1^ Chl min^-1^, respectively; Ku and Edwards, 1975) and *S. anglica* Hubb. (3.6 and 5.7 µmol g^-1^ FW min^-1^; Smith *et al*., 1982; Smith and Woolhouse, 1983). They were not higher and were sometimes lower than in other C_4_ species and C_3_ species. They were lower than the rate of photosynthesis in *M. maximus* (about 6 µmol CO_2_ g^-1^ FW min^-1^ at the growth irradiance of 600 µmol m-^2^ s^-1^, Baccolini *et al*., 2026)

## Discussion

### NAD-ME and PEPK subtypes operate an energy shuttle

It is well-established that NADP-ME species with dimorphic BSC chloroplasts operate an energy shuttle in which part of the 3PGA produced by Rubisco moves from the BSC to the MC where it is reduced to triose-P before returning to the BSC (see Introduction). As NAD-ME and PEPCK species possess granal BSC chloroplasts, the CBC could in principle operate autonomously in the BSC, without exporting 3PGA to the MC. However, indirect evidence including reports of high expression of TPT in the MC and high activities of enzymes that reduce 3PGA to triose-P in MC chloroplasts (see Introduction) suggests NAD-ME and PEPCK species have the capacity to operate an energy shuttle.

Here we provide two lines of experimental evidence that the energy shuttle operates in NAD-ME and PEPCK species. First, a panel of NAD-ME and PEPCK species contained high overall levels of 3PGA and especially DHAP, which is the major triose-P. Levels were comparable to or higher than in NADP-ME species, and almost always higher than in C_3_ species (Figure 1) This extends a previous report of high 3PGA and triose-P in the NAD-ME species *Amaranthus edulis/caudatus* (von Caemmerer and Leegood, 1988). 3PGA and DHAP contents similar to those in the NADP-ME species *Z. mays* and *S. viridis* were also recently reported for the NAD-ME species *P. miliaceum* and the PEPCK species *M. maximus* (Baccolini *et al*., 2026). Second, leaf fractionation revealed that two NAD-ME species (*E. indica, P. miliaceum*) and two PEPCK species (*U. panicoides, M. maximus*) possess a concentration gradient to drive diffusion of 3PGA from the BSC to the MC, and that three (*E. indica, P. miliaceum, U. panicoides*) of the four species possess a concentration gradient to drive diffusion of DHAP from the MC to the BSC (Figure 2). Further, all four species exhibited much higher 3PGA/DHAP ratios in the BSC that the MC (Supplementary Table S6), consistent with energy availability restricting reduction of 3PGA in the BSC and readily available energy driving efficient reduction of 3PGA in the MC.

The estimated amounts and concentrations of 3PGA and DHAP are approximations, for several reasons. First, the fractionation procedure provides only partial enrichment and analysis is complicated by inherent experimental noise in measurements of marker enzyme activities and metabolite levels. Second, the use of marker enzyme distribution to extrapolate to 3PGA and DHAP contents in the BSC and MC depends on assumptions, including that the 3PGA and DHAP are restricted to the BSC and MC. It is likely that small amount is located elsewhere, including other cell types in the vascular material that would be co-enriched with the BSC, and in the epidermic. Third, conversion of metabolite amounts to concentrations depends on assumptions about the cytoplasmic volume of the BSC and MC. Our calculations used published values for *Z. mays* leaves, which may differ from the volumes in the NAD-ME and PEPCK species we investigated. As noted above, there is a high 3PGA/DHAP ratio in the BSC and low ratio in the MC in *M. maximus*. This indicates that *M. maximus* operates an energy shuttle, even though the estimated distribution of DHAP failed to detect a concentration gradient between the MC and BSC. The estimated distribution of 3PGA was very asymmetric for *M. maximus* (Figure S2), which is consistent with the volumes from *Z. mays* overestimating BSC volume and underestimating MC volume in *M. maximus*. This may mask an intercellular gradient for DHAP. This contrasts with the 3PGA/DHAP ratio, which can be compared between the BSC and MC independently of assumptions about cellular volumes.

The estimated intercellular concentration gradients of 3PGA in all four species and of DHAP (excluding *M. maximus*) were of the order of 1.9-7.3 mM and 2.5-6.3mM, respectively (Table 2). The NAD-ME and PEPCK plants were grown and harvested at 550 µmol m^-2^ s^-1^ irradiance. In earlier studies of *Z. mays*, gradients of about 2.9 and 1.4 mM were reported for plants growing at 490 490 µmol m^-2^ s^-1^ irradiance (Arrivault *et al*., 2017) and, and 4 and 9 mM for plants growing at >2000 µmol m^-2^ s^-1^ (Stitt and Heldt 1985a, 1985b) (the published *Z. mays* values have been recalculated with the *Z. mays* volumes used for the calculation in Table 2). The concentration gradient required to drive flux will increase as the rate of photosynthesis and flux around the energy shuttle rises (see Stitt and Heldt, 1985a). Taking this into account, the estimated intercellular gradients in NAD-ME and PEPCK species is in the same range as in the model NADP-ME species *Z. mays*. Furthermore, the range of 3PGA/DHAP ratios in the BSC and MC of the NADP-ME and PEPCK species (2-5-6.7 and 0.35-0.71, respectively) resembles those previously published for *Z. mays*, especially in moderate light (3.7 and 0.68, respectively, Arrivault *et al*., 2017).

### Function of the energy shuttle in NAD-ME and PEPCK species

Overall, our results point to transfer of energy from the MC to the BSC being a general feature of C_4_ photosynthesis, despite the presence of fully functional light reactions in BSC chloroplasts of NAD-ME and PEPCK subtypes. As previously suggested (Hatch, 1987; Edwards and Voznesenskaya, 2011; von Caemmerer and Furbank, 2016), irrespective of whether the BSC chloroplasts possess PSII or not, the energy shuttle allows balancing of energy production and consumption in the MC and BSC. This will include consumption of any ATP and NADPH produced in the MC in excess of that required by the CCM. Furthermore, energy production in the BSC may be somewhat reduced because the BSC is partly shaded by the MC (Bräutigam *et al*., 2014; Stata *et al*., 2014). This might result in an additional need to import energy from the MC in NAD-ME and PEPCK subtypes. Under fluctuating light in canopies there may be rapid changes in shadowing of the BSC. The energy shuttle provides a rapid and flexible way to do this, with transient accumulation of 3PGA in the BSC immediately leading to increased export to the MC. Furthermore, the possibility of energy transfer between MC and BSC will provide flexibility in species that employ multiple decarboxylation pathways, especially as their contribution changes depending on conditions and under fluctuating light (see Medeiros *et al*., 2021; Arrivault *et al*., 2025). It may also, in principle, make it energetically possible to operate a pure PEPCK subtype.

### Why is BSC PSII decreased in NADP-ME species

The existence of 3PGA/triose-P shuttle in NAD-ME and PEPCK species with fully functional BSC chloroplasts raises the following question: did the widespread loss of PSII in NADP-ME species occur because their CCM generates NADPH in the BSC chloroplast and decreases the need for NADPH from PSII, or did it occur in response to other features of the NADP-ME pathway. The finding that an energy shuttle operates in all C_4_ subtypes points to the latter as the more likely explanation. A plausible explanation would be that efficient operation of NADP-ME in the decarboxylation direction to generate a high BSC CO_2_ concentration depends on removal of the other reaction products, especially NADPH. This occurs in the NADPH-GAPDH reaction; but would be comprised if PSII were simultaneously producing NADPH at high rates (Bräutigam *et al*., 2018). It might be noted that an analogous problem does not arise in NAD-ME species, because NAD-ME operates in the mitochondria and the HADH generated by NAD-ME is consumed in the preceding conversion of aspartate to malate by aspartate aminotransferase and NAD-MDH (Hatch, 1987).

### PEP moves back from the BSC to the MC in some PEPK subtypes

The route by which PEP produced by PEPCK moves from the BSC to the MC remains a matter of debate, with possible routes including movement as PEP itself, after conversion to 3PGA or after conversion to pyruvate and alanine (see Introduction). Our study shows that PEP is asymmetrically distributed in two PEPCK species, with larger amounts in the BSC than the MC. The estimated concentration gradient for PEP was 2.2 and 1.1 mM in *M. maximus* and *U. panicoides*, respectively (Table 2), with the larger gradient in *M. maximus* being mainly due to a more asymmetric distribution (Table 1, Figure 2). The concentration gradients are smaller than those for 3PGA (7.3 and 6.4 mM, respectively). On face value, this comparison argues against movement of PEP being the only route but does suggest that some of the PEP returns to the MC as PEP in *M. maximus*, and a smaller part in *U. panicoides*.

However, comparison of the overall magnitude of the PEP and 3PGA concentration gradients may underestimate the relative size of the intercellular concentration gradient for PEP in the cytosol, which is the immediate driving force for diffusion. The concentration gradient in the cytosol depends not only on the overall gradient but also on the distribution of metabolites between the cytosol and chloroplast. Most available knowledge on the subcellular location of metabolites comes from C_3_ plants, but it is unlikely to be fundamentally different in C_4_ plants (see Weiner and Heldt, 1992). Subcellular fractionation experiments (Stitt *et al*., 1980, Gerhardt *et al*. 1987, Weiner and Heldt, 1992; Arrivault *et al*., 2014) showed that 3PGA concentration is higher in the chloroplast than the cytosol whereas DHAP is equally distributed or even slightly higher in the cytosol. This pattern is explained by the chemical properties of 3PGA. The actual substrates for the TPT are 3PGA^2-^ and DHAP^2-^ (Flügge *et al*., 1983). 3PGA has a pK_a_=7.1 and, due to the light-dependent alkalization of the stroma, in the light most of the 3PGA in the chloroplast is present as 3PGA^3-^ that is not transported by TPT (Heldt *et al*., 1973; Gardemann *et al*., 1982; Heldt and Piechulla, 2021). The overall estimated 3PGA concentration gradient therefore probably overestimates the gradient in the cytosol. This may be one reason why the overall gradient for DHAP tends to be smaller than that for 3PGA. It also means that comparison of the overall 3PGA and PEP gradients may underestimate the relative size of the concentration gradient of PEP in the cytosol. This may be further exacerbated by the subcellular distribution of PEP differing from that of 3PGA. Enolase and phosphoglycerate mutase are absent (Stitt and ap Rees, 1979; Bagge and Larsson, 1986) or present at only very low activities (Schulze-Sibert *et al*., 1987) in photosynthetic chloroplasts of C_3_ species. Subcellular fractionation of barley protoplasts revealed that whereas in the cytosol PEP levels were high and in thermodynamic equilibrium with 3PGA, in the chloroplast PEP was very low and strongly displayed from thermodynamic equilibrium with 3PGA (Champigny *et al*., 1992). If the BSC chloroplast resembles a C_3_ chloroplast, PEP will be much higher in the cytosol than in the chloroplast. In the MC this may be different: in species that have PPDK this will produce PEP in the MC chloroplast, and PEP may therefore be lower in the cytosol than the chloroplast. Taken together, the PEP gradient from the BSC to the MC cytosol may be underestimated, and the 3PGA and PEP gradients between the cytosol of BSC and MC could be more similar to each other than indicated by the overall gradients. Furthermore, only part of the decarboxylation in PEPCK species occur via PEPCK, with the rest occurring via NAD-ME or NADP-ME. Taken together, the cytosolic concentration gradient of PEP may be large enough to allow a substantial part of the PEP that is produced by PEPCK in the BSC to move as PEP to the MC in *M. maximus*, and a smaller proportion in *U. panicoides*. This would need to be balanced by movement of Pi from the MC to the BSC, and of amino groups from the BSC to the MC.

If some PEP moves to the MC by another route, especially in *U. panicoides*, which route does it take? Two aspects of our results make it unlikely that it moves as 3PGA. First, the 3PGA/PEP ratio in the BSC (4.0 and 3.2 in) *M. maximus* and *U. panicoides*, respectively) is close to or even above the combined equilibrium constant of the enolase and PGM reactions. Secondly, total enolase and PGM activities in *M. maximus* and *U. panicoides* resembled those in a C_3_ reference species (Supplementary Figure S7), extending earlier findings of Ku and Edwards (1975) for *P. texanum* and *U. panicoides* and Smith *et al*. (1982) and Smith and Woolhouse (1983) for *S. anglica* L. Hubb.. Recent labelling studies in the PEPCK species *M. maximus* (Baccolini *et al*., 2026) also argue against interconversion of PEP and 3PGA making a substantial contribution to movement of PEP to the MC. During^13^ CO_2_ pulse-labelling of *Z. mays*, 3PGA is labelled rapidly but there is a large delay until PEP is labelled (Arrivault *et al*., 2017; Medeiros *et al*., 2022). This delay is explained by relatively slow interconversion of 3PGA and PEP and by rapid flux of unlabeled C into PEP from preexisting pools of pyruvate and alanine (Arrivault *et al*. 2017). Baccholini *et al*. (2026) observed a similar delay in the labelling of PEP in the the PEPCK species *M. maximus* to that in the NADP-ME species *Z. mays* and *S. italicum* and the NAD-ME species *P. miliaceum*. The delay in *M. maximus* is incompatible with movement of PEP to the MC occurring after interconversion with 3PGA as this would lead to rapid mixing of label in 3PGA and PEP. This prompts the question, why interconversion of PEP and 3PGA is relatively slow in C_4_ species, even in PEPCK species where it could in principle facilitate movement of PEP to the MC. One possible explanation would be that effective operation of the CCM and the CBC requires a certain degree of isolation between the two cycles. Taken together, whilst a substantial part of the PEP may return to the MC as PEP in *M. maximus*, the route in *U. panicoides* remains unclear.

In conclusion, a 3PGA/triose-P energy shuttle is a general feature of C_4_ photosynthesis. This will allow flexible and rapid transfer of energy between the MC and BSC not only to balance energy consumption in steady conditions but also during adjustment to changing conditions. The implication is that the abrogation of PSII activity in NADP-ME species is to an adjustment to operation of the energy shuttle but is instead linked to specific requirements to allow efficient decarboxylation by NADP-ME in the BSC chloroplast.

## Author contributions

M.S. and V.C. conceived and planned the experiments. V.C. grew all plants, performed all experiments and analysed enzyme activities, and 3PGA, DHAP and PEP levels, R.F. analysed F6P and FBP. S.A. advised on the leaf fractionation procedure. V.C. prepared the figures and drafted the manuscript, M.S. revised the manuscript and all authors critically reviewed the manuscript before submission.

## Data availability

All data in this study are in the Supplementary Information which are available online.

## Supplementary Material

**Supplementary Figure S1**. Pathway of C_4_ photosynthesis in NADP-ME, NAD-ME and PEPCK subtypes

**Supplementary Figure S2**. Phylogeny of collected C_4_ species

**Supplementary Figure S3**. Fractionation of leaf material by partial homogenization and filtration in liquid N_2_

**Supplementary Figure S4**-Levels of F6P and FBP in selected C_4_ species and the reference C_3_ species *A. thaliana*.

**Supplementary Figure S5**. Contribution of NADP-ME, NAD-ME and PEPCK to the total decarboxylation activity measured in the NADP-ME species *Z. mays*, the NAD-ME specie *E. indica* and *P. miliaceum* and the PEPCK species *M. maximus* and *U. panicoides*.

**Supplementary Figure S6**. Regression analyses of the leaf fractionation experiments to estimate distribution of DHAP and 3PGA between the BSC and MC in selected NAD-ME and PEPCK species.

**Supplementary Figure S7**. In vitro activities of enolase and phosphoglycerate mutase.

**Supplementary Table S1**. Energy requirements of the MC and BSC in different C_4_ subtypes

**Supplementary Table S2**. Species included in the panel of NADP-ME, NAD-ME and PEPCK subtypes

**Supplementary Table S3**. Overall content of 3PGA and DHAP.

**Supplementary Table S4**. Overall content of F6P and FBP

**Supplementary Table S5**. Estimated distribution of 3PGA and DHAP between the MC and BSC in selected NAD-ME and PEPCK species.

**Supplementary Table S6**. 3PGA/DHAP ratio in the BSC and MC in selected NAD-ME and PEPCK species.

**Supplementary Table S7**. Estimated distribution of PEP between the MC and BSC in the PEPCK subtypes *M. maximus* and *U. panicoides* (supplementary to Figure 2).

**Supplementary Dataset S1**. Measurements of marker enzymes, 3PGA, DHAP and PEP in material from liquid N_2_ fractionation of leaves

